# Linking Gene Expression to Clinical Outcomes in Pediatric Crohn’s Disease Using Machine Learning

**DOI:** 10.1101/2022.11.07.515480

**Authors:** Kevin A Chen, Nina Nishiyama, Meaghan M Kennedy Ng, Alexandra Shumway, Chinmaya U Joisa, Matthew R Schaner, Grace Lian, Caroline Beasley, Lee-Ching Zhu, Surekha Bantumilli, Muneera R Kapadia, Shawn M Gomez, Terrence S Furey, Shehzad Z Sheikh

**Author notes:** Correspondence should be addressed to TSF and SZS.

## Abstract

**Introduction:** Pediatric Crohn’s disease (CD) is the fastest growing age group and is characterized by frequent disease complications. We sought to analyze both ileal and colonic gene expression in a cohort of pediatric CD patients and apply machine learning-based models to predict risk of developing future complications.

**Methods:** RNA-seq was generated from matched ileal and colonic biopsies from formalin-fixed, paraffin-embedded (FFPE) tissue obtained from patients with non-stricturing/non-penetrating, treatment-naïve CD and from controls. Clinical outcomes including development of strictures or fistulas, progression to surgery, and remission were analyzed first using differential expression. Machine learning models were then developed for each outcome, combining gene expression and clinical factors. Models were assessed using area under the receiver operating characteristic curve (AUROC).

**Results:** 56 patients with CD and 46 controls were included. Differential expression analysis revealed a distinct colonic transcriptome for patients who developed strictures, with downregulation of pathways related to inflammation and extra-cellular matrix production. In contrast, there were few differentially expressed genes for other outcomes and for ileal tissue. Despite this, machine learning-based models were able to incorporate colonic gene expression and clinical characteristics to predict outcomes with high accuracy. Models showed an AUROC of 0.84 for strictures, 0.83 for remission, and 0.75 for surgery. Certain genes with potential prognostic importance for strictures (REG1A, MMP3, and DUOX2) were not identified in single gene differential analysis but were found to have strong contributions to predictive models.

**Conclusions:** Our findings in FFPE tissue support the importance of colonic gene expression and the potential for machine learning-based models in predicting outcomes for pediatric CD.

## Introduction

Pediatric Crohn’s disease (CD) is the fastest growing incident age group for the disease with about 80,000 children in the US affected.^1–3^ CD is characterized by a relapsing, remitting disease course with complications, such as strictures or perforation, affecting around 50% of patients within 5 years of diagnosis.^4,5^ Pediatric CD follows a more severe disease course, more often involving strictures and fistulas.^6–8^ These complications drive further morbidity and healthcare utilization associated with CD including growth failure, delayed puberty, hospitalizations, and surgery.^4,8^

Analysis of gene expression and identification of biological pathways which drive development of CD and CD complications may give insight into more precise treatment decision-making to prevent a complicated CD course. Genes associated with immune and cytokine pathways have been associated with CD development.^9–11^ Further, specific genes including oncostatin M, IL1B, S100A8, and CXCL1 have been associated with response to anti-tumor necrosis factor therapy.^12,13^ Genes controlling extracellular matrix production and inflammatory processes have been associated with strictures.^14–16^ Decision-support tools which incorporate this genetic information to prognosticate disease course could assist with clinical decision-making.

Multiple previous studies have sought to predict outcomes for CD based on gene expression, most notably using the RISK cohort.^14^ However, these studies relied on logistic regression models, which may fail to capture the multi-factorial, non-linear interactions between genes and clinical characteristics that connote increased risk for complications. Machine learning techniques, which have the capacity to capture these complex patterns, have been successfully applied to inflammatory bowel disease (IBD)-related topics including identification of risk genes, prediction of outcomes from serum proteins, and prediction of response to medication from multi-omic data.^17–19^ However, they have not yet been applied specifically to prediction of complications for pediatric CD from gene expression.

The goals of our study are twofold: to identify genes which are differentially expressed in CD and complicated CD and to apply machine learning techniques that use those genes to predict risk of complications. We hypothesize that machine learning techniques can incorporate the gene expression profiles of patients with complicated disease to outperform previous predictors.

## Methods

### Study design and outcomes

This study included patient data that was collected at the University of North Carolina at Chapel Hill.^20^ This included patients younger than 18 with suspected IBD, who underwent endoscopy between 2008 and 2012. Patients who were found to have no gut inflammation were used as non-IBD controls. At the time of diagnosis, patients were selected based on non-penetrating, non-stricturing disease phenotype. Parents or guardians of all patients provided written consent and patients provided assent when appropriate. This study was approved by the University of North Carolina Institutional Review Board (Study ID#: 15-0024, 11-0359, 17-0236).

Disease behavior was defined according to the Montreal classification system. Disease complications included strictures (B2), fistulas (B3), progression to surgery, and experiencing remission. B2 and B3 disease were defined using endoscopy and/or imaging (fluoroscopy, CT, or MRI) and correlation with patient symptoms in contrast to the non-stricturing, non-fistulizing phenotype (B1).^21,22^ Progression to surgery was defined as requiring an abdominal surgical procedure for resection of bowel. Remission was defined as experiencing a steroid-free interval of at least 6 months.^9^ Outcomes were recorded with a mean follow-up period of 6 years.

### Specimen, mRNA, and data processing

Macroscopically uninflamed mucosal samples from the ascending colon and terminal ileum were obtained at the time of initial diagnosis, before therapy was started. These samples were preserved as fresh frozen paraffin-embedded (FFPE) tissue.

RNA was isolated from FFPE tissue using the Quick-RNA FFPE MiniPrep (Zymo Research, Irvine, CA). This kit preserves mRNA content while using column-based DNase to eliminate DNA contamination. Total RNA was then purified using the MagMAX kit in the KingFisher system (ThermoFisher, Carlsbad, CA). RNA-seq libraries were prepared using TruSeq Stranded Total RNA with Ribo-Zero (Illumina, San Diego, CA). Paired-end (50bp) sequencing was processed on the NovaSeq 6000 platform using default parameters (Illumina, San Diego, CA). Transcript expression was then quantified using Salmon with default parameters.^23^ Purity and integrity of the samples was assessed using a variety of quality control metrics. We first identified samples with a low number of transcripts counted (<25,000). Further investigation of these samples confirmed low transcript integrity number (TIN),^24^ percentage of sequences aligned, and high duplication percentage. These samples (n = 2) were then discarded. Further, we used PCA (principal component analysis) plots to identify samples which did not cluster with their respective tissue (ileal or colonic) and discarded these samples as well (n = 5).

### Statistical analysis

PCA showed that batch, sex, and TIN drove the greatest variation between samples that was unrelated to disease phenotype, so these variables were explicitly included as covariates. Additional factors of unwanted variation were identified using RUVSeq.^25^ Control genes were selected by identifying the top 1000 genes with the lowest variance out of the top 5000 genes with the highest expression. Based on variation seen in relative log expression plots across samples, correlation between factors of unwanted variation and the outcome, and the number of differentially expressed genes identified by DESeq2, we used one factor of unwanted variation for final analyses.

Final PCA plots were generated using the plotPCA function from DESeq2, based on the top 500 most variable genes, after applying the variance stabilizing transform (VST) and the removeBatchEffect function from limma.^26,27^ The filterbyExpression function from EdgeR was used to select genes with at least 10 read counts in 70% of samples.^28^ Differential expression analysis was then performed using DESeq2 with false discovery rate (FDR) adjusted P-value (p-adj) of <0.05 considered significant. Pathway analysis was performed using the Molecular Signatures Database hallmark gene set collection and fgsea.^29,30^ Volcano plots were generated using EnhancedVolcano.^31^ RNA-seq analysis was performed in R (v4.2).^32^

### Modeling

Predictive models were developed for the collected outcomes, including development of B2 phenotype, progression to surgery, and remission. Consecutive models were built including clinical variables alone (Table 1) and clinical variables with gene expression in order to evaluate the contribution of gene expression to overall predictions. Separate models were also built with and without rectosigmoid involvement, a clinical feature not previously reported in other predictive models for pediatric CD.^20,33^ Based on the results of the differential expression analysis, colonic gene expression data was used. Models were trained based on normalized gene counts, processed as described above including filtering genes by expression, controlling for batch, sex, TIN, and 1 factor of variation, and normalizing using the variance stabilizing transform.^25,26,28^ Given the small sample size, leave-one-count cross-validation was used. With this approach, a unique model is trained for each sample in the dataset, that sample is excluded from training and used for evaluation, and model performance is represented as an average across all samples. Genes were selected for inclusion within models using the least absolute shrinkage and selection operator (LASSO), a regularized linear model that increases a penalty for each non-zero coefficient.^34^ Care was taken to apply gene selection within folds, with LASSO applied to only the training data for each fold.

**Table 1.**
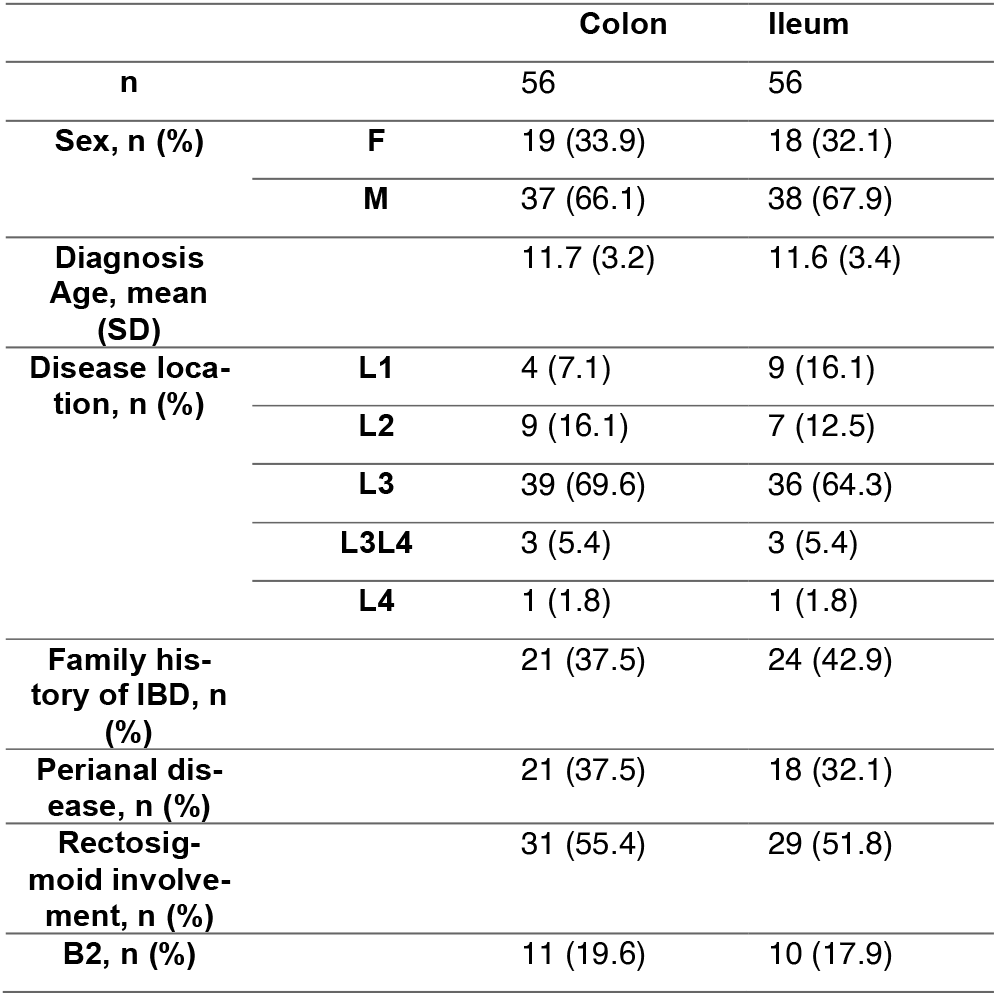
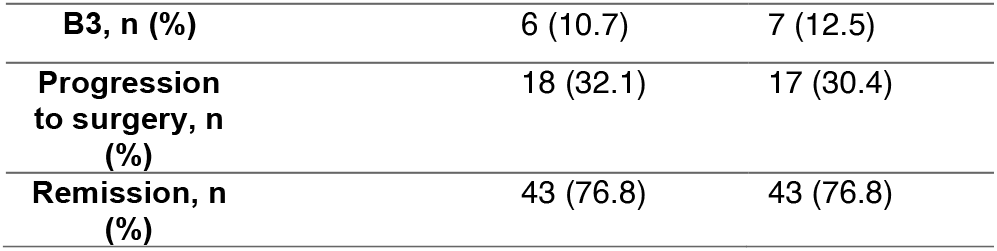
Clinical and Demographic Characteristics of the Crohn’s Disease Study Cohort.

|  |  | Colon | Ileum |
| --- | --- | --- | --- |
| **n** |  | 56 | 56 |
| **Sex, n (%)** | **F** | 19 (33.9) | 18 (32.1) |
|  | **M** | 37 (66.1) | 38 (67.9) |
| **Diagnosis Age, mean (SD)** |  | 11.7 (3.2) | 11.6 (3.4) |
| **Disease location, n (%)** | **L1** | 4 (7.1) | 9 (16.1) |
|  | **L2** | 9 (16.1) | 7 (12.5) |
|  | **L3** | 39 (69.6) | 36 (64.3) |
|  | **L3L4** | 3 (5.4) | 3 (5.4) |
|  | **L4** | 1 (1.8) | 1 (1.8) |
| **Family history of IBD, n (%)** |  | 21 (37.5) | 24 (42.9) |
| **Perianal disease, n (%)** |  | 21 (37.5) | 18 (32.1) |
| **Rectosigmoid involvement, n (%)** |  | 31 (55.4) | 29 (51.8) |
| **B2, n (%)** |  | 11 (19.6) | 10 (17.9) |

|  |  |  |
| --- | --- | --- |
| **B3, n (%)** | 6 (10.7) | 7 (12.5) |
| **Progression to surgery, n (%)** | 18 (32.1) | 17 (30.4) |
| **Remission, n (%)** | 43 (76.8) | 43 (76.8) |

Multiple machine learning models were developed and compared, including LASSO, random forest (RF), gradient boosting (XGB), deep neural networks (NN).^35^ Each model was assessed using area under the receiver operating characteristic curve (AUROC) and area under the precision-recall curve (AUPRC). Feature importance was determined for the LASSO model using its coefficients. Coefficients were summarized across cross-validation folds by summing the absolute value for each fold. PCA plots were then generated using the genes with the highest coefficient values across all folds. Model training, evaluation, and interpretation was performed in Python (v3.8) using the Scikit-Learn and Tensorflow libraries.^35–37^

## Results

### Study population characteristics

After applying quality control, 56 CD patients with colon samples and 56 CD patients with ileum samples were included in the study cohort, while 46 non-IBD patients with colon samples and 46 non-IBD patients with ileum samples were used as controls. For CD patients with colon samples, 33.9% of patients were female, the average age of diagnosis was 11.7, and 69.6% of patients had ileocolonic disease.

19.6% of patients developed B2 complications, 10.7% developed B3 complications, 32.1% required surgery, and 76.8% experienced a period of remission (Table 1). Of note, all 12 patients who developed B2 complications required surgery and 12 of 19 (63.1%) of patients who required surgery had B2 complications.

### Differential expression analysis

We first identified differentially expressed genes (DEG’s) between patients with CD compared with non-IBD controls, in both colonic and ileal tissue. In total, 10,973 DEG’s were identified for colonic tissue and 8,799 for ileal tissue (p-adj < 0.05) (Figure 1C/D). Genes related to inflammatory response (CXCL8, AQP9, INHBA, IL1B, CXCL6, and IL6) were upregulated in CD compared with non-IBD, while genes related to DNA repair (MPC2, VPS28, EDF1, ALYREF, and PCNA) and oxidative phosphorylation (IDH3B, ATP5MC1, ATP5ME, MRPL11, COX7C, and PHB2) were downregulated. A complete list of all differential expression results is available in Supplementary Table 1 (colon) and 2 (ileum).

**Figure 1.**
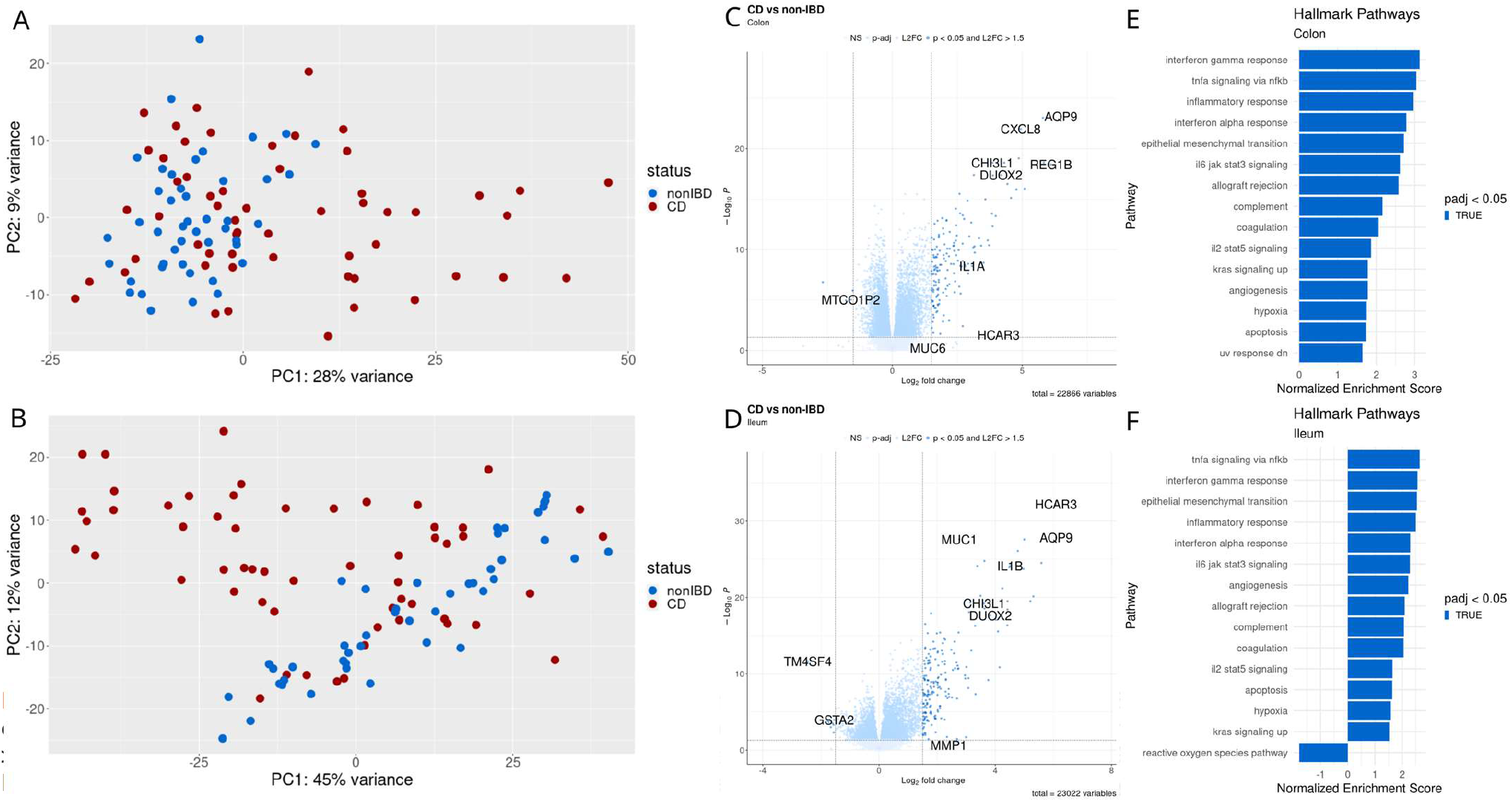
Differential gene expression analysis for pediatric patients with CD versus patients without IBD. A, PCA plot based on colonic gene expression. B, PCA plot based on ileal gene expression. C, Volcano plot showing differentially expressed genes with p < 0.05 and log2 fold change > 1.5 based on colonic gene expression. D, Volcano plot based on ileal gene expression (same criteria). E, Gene set enrichment analysis based on Hallmark pathways for colonic gene expression. F, Gene set enrichment analysis based on Hallmark pathways for ileal gene expression.

We then analyzed DEG’s between patients experiencing specific outcomes (B2 – stricturing, B3 – fistulizing, progression to surgery, and remission) and those who did not. Of the four outcomes, B2 showed the clearest difference in gene expression. For colonic tissue, genes related to extra-cellular matrix (ECM) production (MMP3, MMP1, CHI3L1), as well as inflammatory processes (CXCL5, CXCL8, AQP9, INHBA) were downregulated in patients who experienced B2 complications. The Hallmark pathways interferongamma response, inflammatory response, and epithelial mesenchymal transition were notably downregulated. A full list of differential expression results for B2 in colonic tissue is available in Supplementary Table 3. For B2 in ileal tissue, no significantly DEG’s were identified. Analysis of DEG’s for B3 showed 2 for colon and 1 for ileum, although these showed no specific pattern. For progression to surgery, 4 DEG’s were identified for colon and 1 for ileum. This included upregulation of mitochondrial genes (MTCO1P12 and MTND1P23) and downregulation of UCN2 and CXCL5 in colonic tissue. For ileal tissue, MTCO1P12 was upregulated. Finally, analysis of remission showed no DEG’s.

### Predictive modeling

We first developed models for each of the recorded outcomes based on clinical variables alone (sex, diagnosis age, disease location, perianal disease, and family history of IBD). Overall, these showed poor accuracy with AUROC of <0.6 for all models for all outcomes. Adding gene expression resulted in a significant improvement in predictive ability (Figure 3). For B2, neural networks (NN) showed the highest performance, with an AUROC of 0.806 (95% CI 0.753 - 0.859) compared with 0.583 (95% CI 0.518 - 0.649) for clinical variables alone. For remission and surgery, NN was the highest performing model, obtaining an AUROC of 0.834 (95% CI 0.784 - 0.883) and 0.732 (95% CI 0.673 - 0.792) for each outcome respectively. AUROC and AUPRC results for all models are available in Supplementary Table 4.

**Figure 2.**
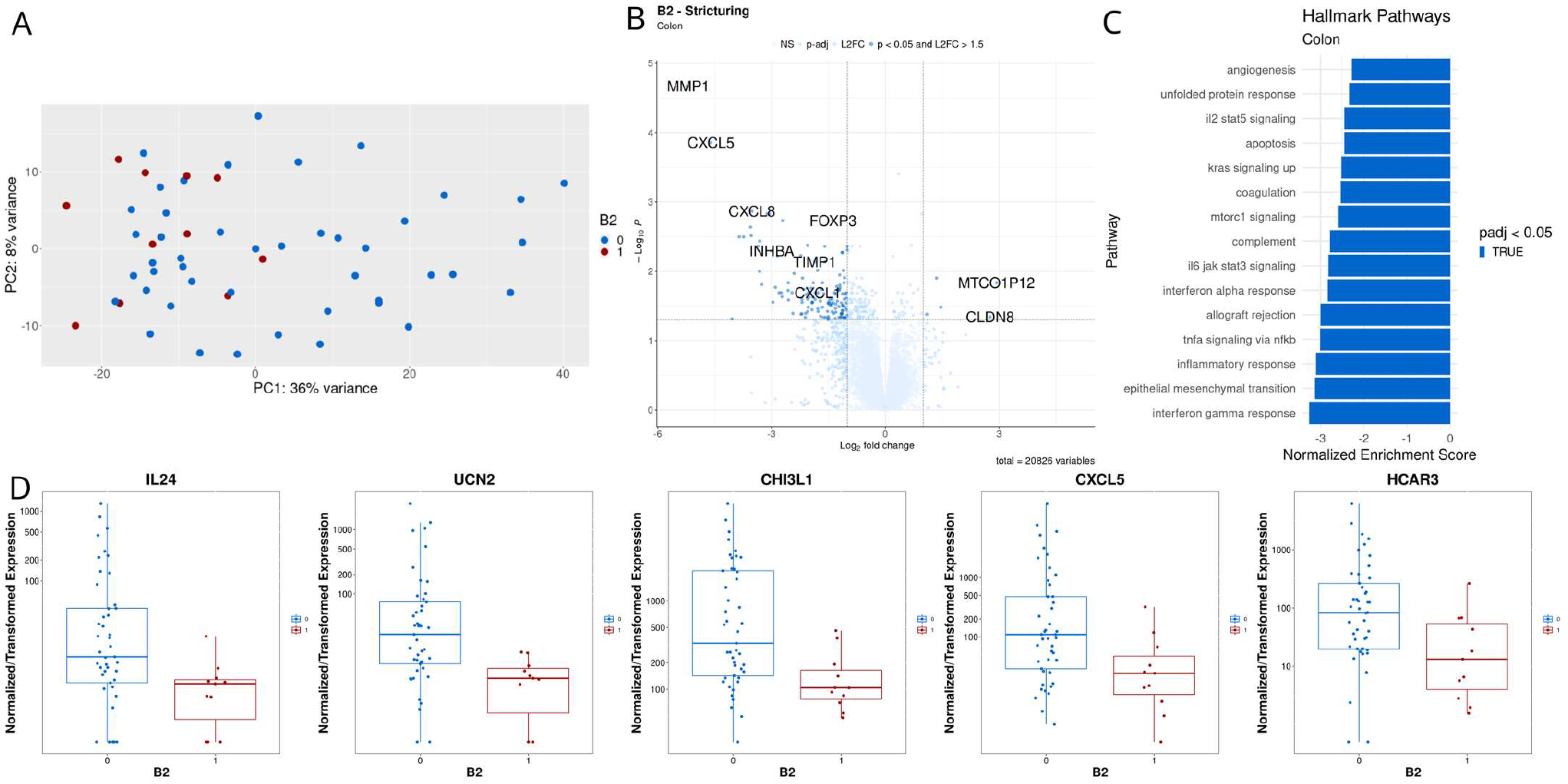
Differential gene expression analysis for pediatric CD patients experiencing stricturing complications versus those who did not based on colonic tissue. A, PCA plot. B, Volcano plot showing differentially expressed genes with p < 0.05 and log2 fold change > 1.5. C, Gene set enrichment analysis based on Hallmark pathways. D, Boxplots for selected genes

**Figure 3.**
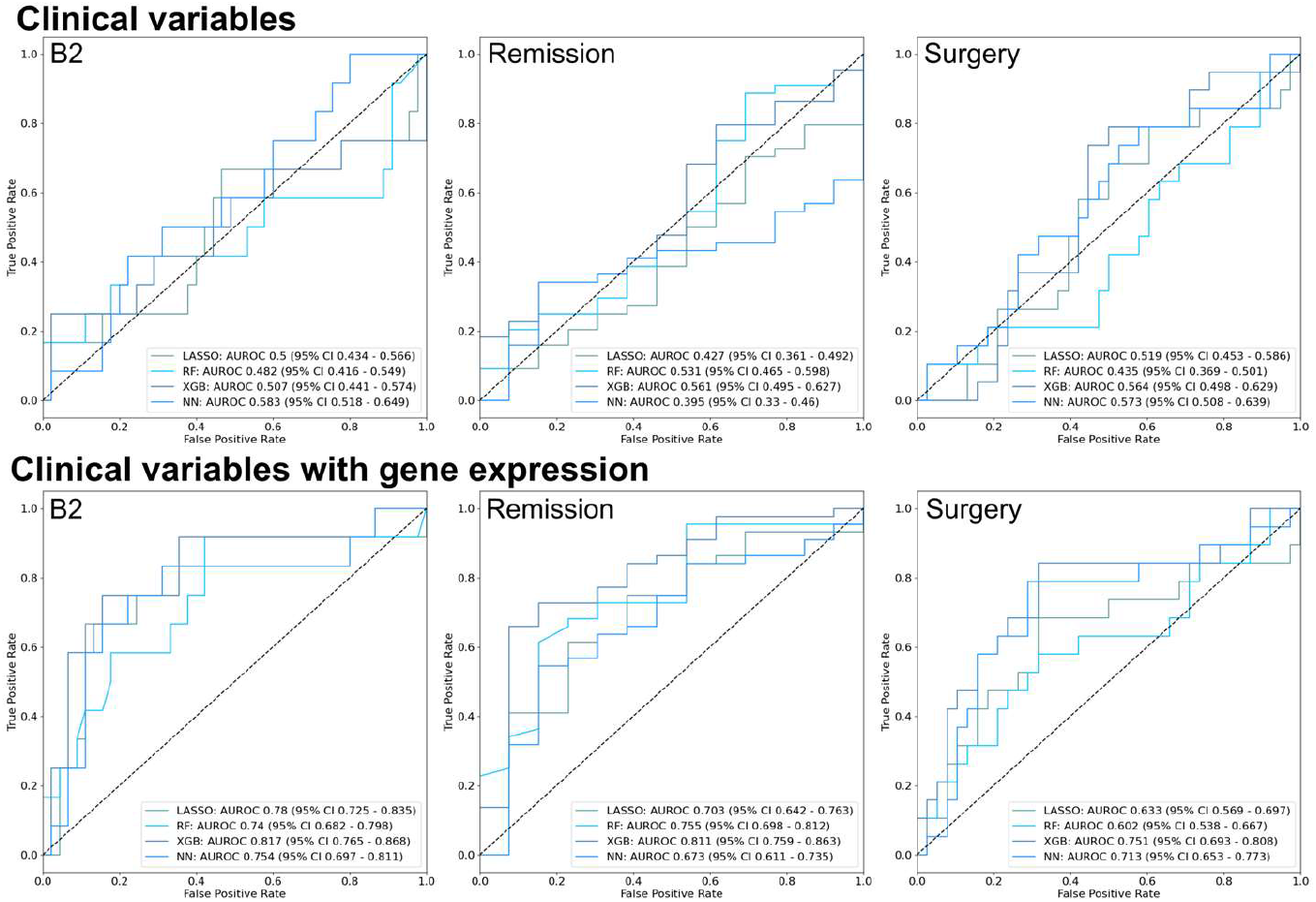
Receiver operating characteristic curves for all models predicting pediatric CD complications based on clinical variables and gene expression, RF – random forest, XGB – gradient boosting, NN – neural network, AUROC - area under the receiver operating characteristic curve, CI – confidence interval

Addition of rectosigmoid involvement to the clinical model also resulted in significant improvements for all outcomes compared the original clinical variables with AUROC 0.7-0.8. Finally, combining all variable types (clinical variables, rectosigmoid involvement, and gene expression) resulted in the highest accuracy for B2, with NN showing an AUROC of 0.836, and remission, with XGB showing an AUROC of 0.834 (Figure 4). In contrast, for surgery, clinical variables with gene expression and clinical variables with rectosigmoid involvement showed the best performance, with an AUROC for gradient boosting (XGB) of 0.751. AU-ROC and AUPRC results for these models are available in Supplementary Table 4. Analysis of the LASSO prediction model for B2 to determine which genes showed the strongest contributions to model predictions revealed differences compared with differential expression analysis. Of the 131 genes used across all folds, 33 were found to be significantly differentially expressed. Genes related to inflammatory/immune processes were highly important, including CXCL9, DUOX2, and FOXP3. ECM-related genes were also important, including MMP3, MMP1, and CHI3L1. Genes with the largest cumulative absolute values for coefficients are listed in Figure 5A. Pathway enrichment analysis showed that the Hallmark pathways interferon-gamma response and IL-6/JAK/STAT signaling showed the strongest enrichment (Figure 5B). PCA plots based only on the top 20 genes identified by the LASSO models showed strong clustering of the B2 samples (Figure 5C). Interestingly, of the 5 genes used in >50% of folds (REG1A, FGL2, DMBT1, MMP3, and DUOX2), only 1 (DMBT1) was found to be significantly differentially expressed. Two of these, FGL2 and DUOX2 trended towards significance, with adjusted p-values of 0.17 and 0.07 respectively. Analysis of expression of these specific genes showed clear differences between the two groups, but significant heterogeneity.

**Figure 4.**
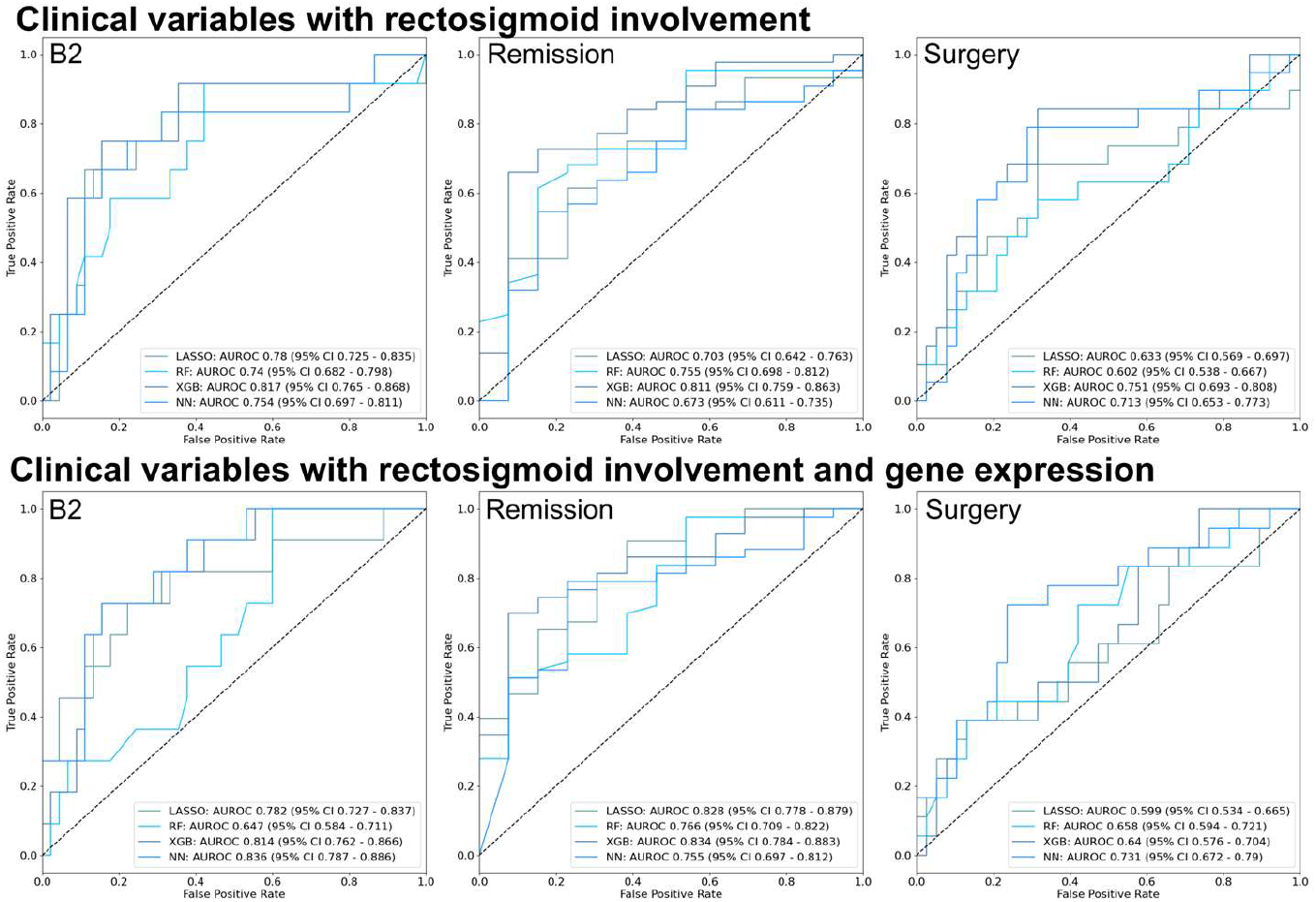
Receiver operating characteristic curves for all models predicting pediatric CD complications based on clinical variables, rectosigmoid involvement, and gene expression, RF – random forest, XGB – gradient boosting, NN – neural network, AUROC - area under the receiver operating characteristic curve, CI – confidence interval

**Figure 5.**
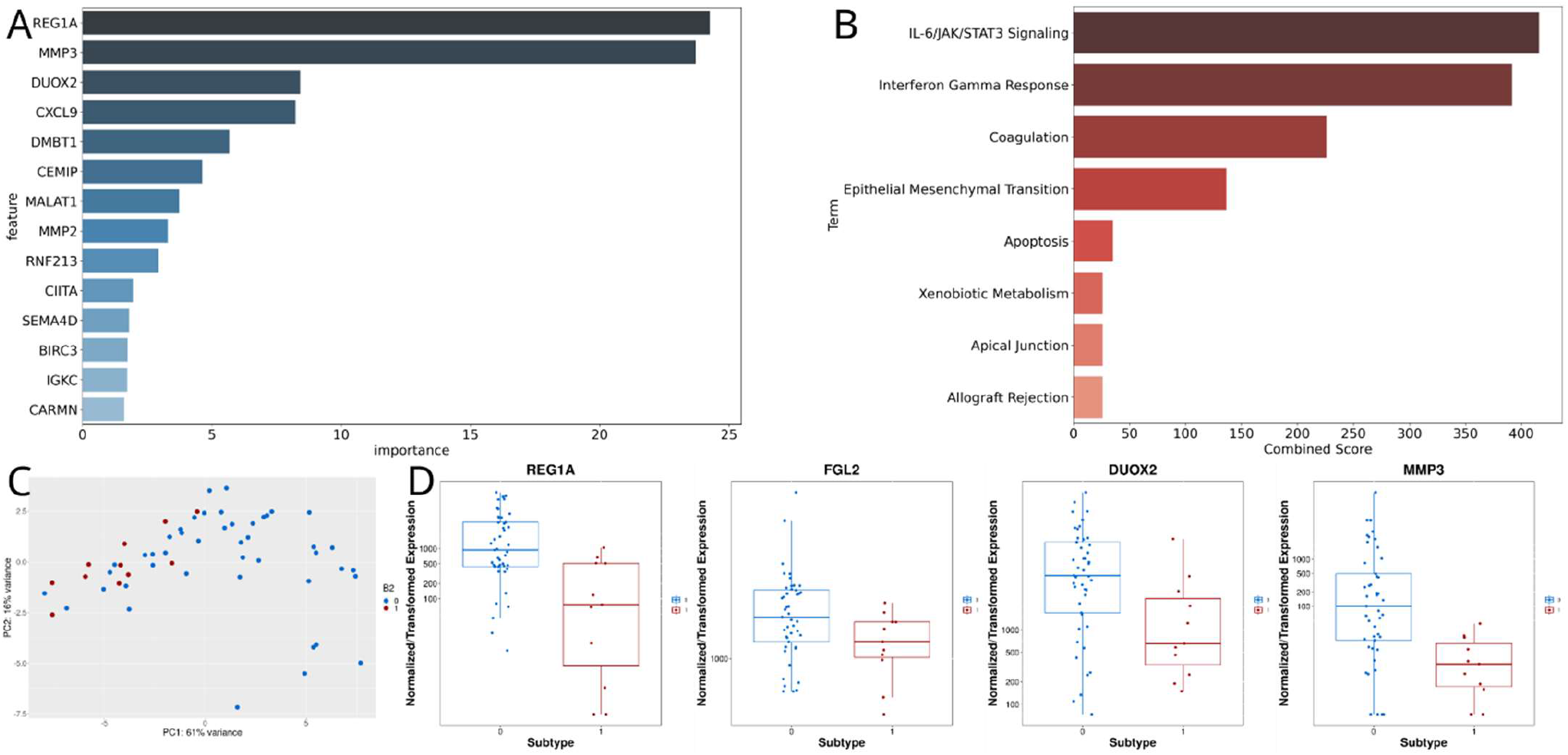
Analysis of model predicting stricturing (B2) complications for pediatric CD. A, Top genes based on LASSO coefficients across all cross-validation folds. B, Pathway analysis based on top genes. C, PCA plot based on top genes. D, Boxplots of expression by B2 status for genes used in >50% of folds, but not found to be differentially expressed

## Discussion

Patients with pediatric CD who experienced stricturing complications showed a distinct colonic transcriptome at time of diagnosis compared with those who did not, with downregulation of inflammatory and extracellular matrix (ECM) production pathways. Patients who required surgery also showed downregulation of the ECM-related pathways. In contrast, there was no clear difference in the pattern of gene expression between patients who experienced fistulizing complications or those who experienced remission based on differential expression analysis. Machine learning-based models were able to incorporate information from gene expression to improve upon predictions based on clinical variables alone and predict with high accuracy which patients would develop stricturing complications, experience remission, or require surgery. Despite limited changes in individual genes for the remission and surgery outcomes, the models were able to achieve good accuracy, suggesting improved predictions based on combinations of genes.

Multiple previous studies have established a link between gene expression, particularly in the ECM and inflammatory pathways, and pediatric CD outcomes.^38^ Haberman et al. identified increased *DUOX2, MMP3, AQP9*, and *IL8* as highly upregulated and *APOA1, NAT8*, and *AGXT2* as highly downregulated in ileal tissue for pediatric CD. These gene signatures were then used to predict steroid-free remission with an AUROC of 0.721.^9^ Kugathasan et al. identified upregulation of several ECM-related gene ontology pathways in the ileum of pediatric CD patients experiencing B2 complications and used an ECM gene signature to predict development of B2 complications with an AUROC of 0.72.^14^ Ta et al. also identified inflammatory and ECM gene signatures as associated with transmural healing for pediatric CD patients with inflammatory small bowel disease.^39^

The results of our study broadly agree with previous work and confirm the importance of ECM and inflammatory pathways for pediatric CD outcomes. However, they also differ from previous work in pediatric CD in that our analysis focuses on colonic rather than ileal tissue and shows downregulation of the inflammatory response and epithelial mesenchymal transition pathways in this tissue type. The current results agree with previous studies suggesting prognostic significance of colonic gene expression for predicting mainly ileal complications, as the ileal transcriptome may be completely dominated by current, active disease.^21,40^ Of note, these results relied on FFPE tissue, which allowed assembly of a broader cohort at lower cost, but showed broad agreement with results based on fresh tissue, especially in CD vs non-IBD comparisons. In addition, despite using a smaller training set and rigorous cross-validation, our models show higher predictive accuracy (AUROC >0.8) compared with previous studies, demonstrating the potential for more complex, machine learning-based models to outperform traditional logistic regression.

Analysis of the contributions of individual genes to our models reveals associations between genes and outcomes that may be overlooked by single gene differential expression techniques. Due to heterogeneity in gene expression, these associations may not appear when groups are considered in aggregate. In particular, the genes *REG1A, MMP3*, and *DUOX2* strongly influenced model predictions and have been found to be associated with IBD and disease severity in multiple previous studies, but were not identified as significantly differentially expressed.^9,41,42^ Another interesting finding from our study was the strong inverse relationship between rectosigmoid involvement and development of stricturing disease. Previous studies have identified young age, ileocolonic involvement, perianal involvement, and early response to initial therapy as predictive of CD complications.^5,33,43^ However, few studies have specifically examined rectosigmoid disease.^43^ This finding merits further study in other populations.

Our results join a growing body of research highlighting the potential for machine learning to predict outcomes related to IBD and support clinicians in providing therapies tailored to those predictions. Machine learning has been used to predict hospitalization and outpatient steroid use,^44^ response to biologic therapy,^45^ post-operative CD recurrence,^46^ and identify novel serum markers.^47^ Machine learning can identify relationships within multi-omic, high dimensional data and is particularly well-suited to assist the transition from a “trial and error” approach to precision medicine in IBD.^48^

Our study has important limitations. First, it is based on a relatively small, single-institution dataset. While the exact models generated using this dataset may not be generalizable, the described methods for selecting and modeling on gene expression should be broadly applicable. Second, similar to previous studies, we were not able to consistently model B3 complications, likely due to the heterogeneity of the subtype.^14^ Third, analyzing paired affected and unaffected regions for each patient may have captured the impact of inflammation on molecular phenotypes. Fourth, treatment in this study was left to the discretion of the primary pediatric gastroenterologist and differences in treatment selection had an unadjusted effect on outcomes. Finally, our analysis does not include other data types, such as small RNA, chromatin biology, serum markers, or microbial composition. Prediction of IBD outcomes applying machine learning to these multi-omic data sources represents an exciting direction for future research.^19,49^

## Conclusions

Pediatric CD patients who experience complications show a distinct colonic transcriptome at the time of diagnosis. Machine learning can use this information to predict future outcomes, including strictures, remission, or progression to surgery. Applied to larger, multi-institutional datasets, this approach can develop prognostic models to support clinicians in identifying which patients are at highest risk of CD-specific complications and tailor therapies to improve outcomes.

## Acknowledgements

This study was supported by work from the University of North Carolina Translational Pathology Lab, High Throughput Sequencing Facility, and Tissue Genomic Lab which are supported in part by an NCI Center Core Support Grant (5P30CA016080-42).

This paper was typeset with the bioRxiv word template by @Chrelli: www.github.com/chrelli/bioRxiv-word-template

## Supplementary Content

**Supplementary Table 1.**
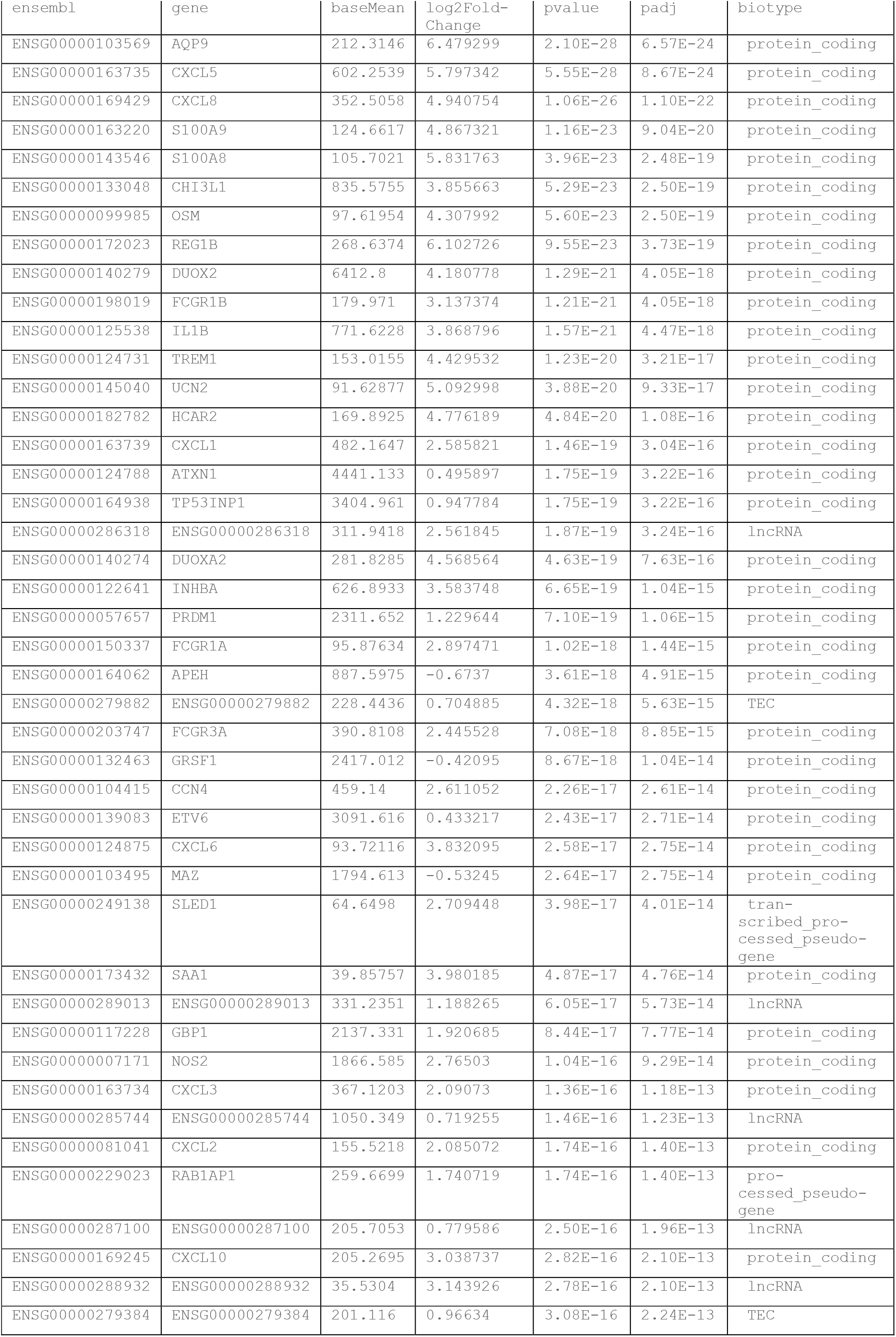
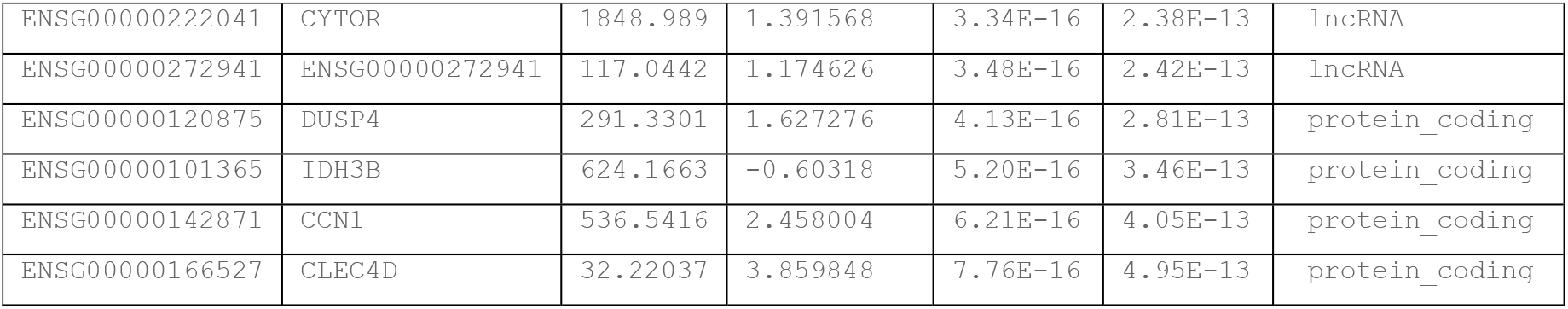

| ensembl | gene | baseMean | log2Fold-Change | pvalue | padj | biotype |
| --- | --- | --- | --- | --- | --- | --- |
| ENSG00000103569 | AQP9 | 212.3146 | 6.479299 | 2.10E-28 | 6.57E-24 | protein_coding |
| ENSG00000163735 | CXCL5 | 602.2539 | 5.797342 | 5.55E-28 | 8.67E-24 | protein_coding |
| ENSG00000169429 | CXCL8 | 352.5058 | 4.940754 | 1.06E-26 | 1.10E-22 | protein_coding |
| ENSG00000163220 | S100A9 | 124.6617 | 4.867321 | 1.16E-23 | 9.04E-20 | protein_coding |
| ENSG00000143546 | S100A8 | 105.7021 | 5.831763 | 3.96E-23 | 2.48E-19 | protein_coding |
| ENSG00000133048 | CHI3L1 | 835.5755 | 3.855663 | 5.29E-23 | 2.50E-19 | protein_coding |
| ENSG00000099985 | OSM | 97.61954 | 4.307992 | 5.60E-23 | 2.50E-19 | protein_coding |
| ENSG00000172023 | REG1B | 268.6374 | 6.102726 | 9.55E-23 | 3.73E-19 | protein_coding |
| ENSG00000140279 | DUOX2 | 6412.8 | 4.180778 | 1.29E-21 | 4.05E-18 | protein_coding |
| ENSG00000198019 | FCGR1B | 179.971 | 3.137374 | 1.21E-21 | 4.05E-18 | protein_coding |
| ENSG00000125538 | IL1B | 771.6228 | 3.868796 | 1.57E-21 | 4.47E-18 | protein_coding |
| ENSG00000124731 | TREM1 | 153.0155 | 4.429532 | 1.23E-20 | 3.21E-17 | protein_coding |
| ENSG00000145040 | UCN2 | 91.62877 | 5.092998 | 3.88E-20 | 9.33E-17 | protein_coding |
| ENSG00000182782 | HCAR2 | 169.8925 | 4.776189 | 4.84E-20 | 1.08E-16 | protein_coding |
| ENSG00000163739 | CXCL1 | 482.1647 | 2.585821 | 1.46E-19 | 3.04E-16 | protein_coding |
| ENSG00000124788 | ATXN1 | 4441.133 | 0.495897 | 1.75E-19 | 3.22E-16 | protein_coding |
| ENSG00000164938 | TP53INP1 | 3404.961 | 0.947784 | 1.75E-19 | 3.22E-16 | protein_coding |
| ENSG00000286318 | ENSG00000286318 | 311.9418 | 2.561845 | 1.87E-19 | 3.24E-16 | lncRNA |
| ENSG00000140274 | DUOXA2 | 281.8285 | 4.568564 | 4.63E-19 | 7.63E-16 | protein_coding |
| ENSG00000122641 | INHBA | 626.8933 | 3.583748 | 6.65E-19 | 1.04E-15 | protein_coding |
| ENSG00000057657 | PRDM1 | 2311.652 | 1.229644 | 7.10E-19 | 1.06E-15 | protein_coding |
| ENSG00000150337 | FCGR1A | 95.87634 | 2.897471 | 1.02E-18 | 1.44E-15 | protein_coding |
| ENSG00000164062 | APEH | 887.5975 | -0.6737 | 3.61E-18 | 4.91E-15 | protein_coding |
| ENSG00000279882 | ENSG00000279882 | 228.4436 | 0.704885 | 4.32E-18 | 5.63E-15 | TEC |
| ENSG00000203747 | FCGR3A | 390.8108 | 2.445528 | 7.08E-18 | 8.85E-15 | protein_coding |
| ENSG00000132463 | GRSF1 | 2417.012 | -0.42095 | 8.67E-18 | 1.04E-14 | protein_coding |
| ENSG00000104415 | CCN4 | 459.14 | 2.611052 | 2.26E-17 | 2.61E-14 | protein_coding |
| ENSG00000139083 | ETV6 | 3091.616 | 0.433217 | 2.43E-17 | 2.71E-14 | protein_coding |
| ENSG00000124875 | CXCL6 | 93.72116 | 3.832095 | 2.58E-17 | 2.75E-14 | protein_coding |
| ENSG00000103495 | MAZ | 1794.613 | -0.53245 | 2.64E-17 | 2.75E-14 | protein_coding |
| ENSG00000249138 | SLED1 | 64.6498 | 2.709448 | 3.98E-17 | 4.01E-14 | transcribed_processed_pseudo-gene |
| ENSG00000173432 | SAA1 | 39.85757 | 3.980185 | 4.87E-17 | 4.76E-14 | protein_coding |
| ENSG00000289013 | ENSG00000289013 | 331.2351 | 1.188265 | 6.05E-17 | 5.73E-14 | lncRNA |
| ENSG00000117228 | GBP1 | 2137.331 | 1.920685 | 8.44E-17 | 7.77E-14 | protein_coding |
| ENSG00000007171 | NOS2 | 1866.585 | 2.76503 | 1.04E-16 | 9.29E-14 | protein_coding |
| ENSG00000163734 | CXCL3 | 367.1203 | 2.09073 | 1.36E-16 | 1.18E-13 | protein_coding |
| ENSG00000285744 | ENSG00000285744 | 1050.349 | 0.719255 | 1.46E-16 | 1.23E-13 | lncRNA |
| ENSG00000081041 | CXCL2 | 155.5218 | 2.085072 | 1.74E-16 | 1.40E-13 | protein_coding |
| ENSG00000229023 | RAB1AP1 | 259.6699 | 1.740719 | 1.74E-16 | 1.40E-13 | processed_pseudo-gene |
| ENSG00000287100 | ENSG00000287100 | 205.7053 | 0.779586 | 2.50E-16 | 1.96E-13 | lncRNA |
| ENSG00000169245 | CXCL10 | 205.2695 | 3.038737 | 2.82E-16 | 2.10E-13 | protein_coding |
| ENSG00000288932 | ENSG00000288932 | 35.5304 | 3.143926 | 2.78E-16 | 2.10E-13 | lncRNA |
| ENSG00000279384 | ENSG00000279384 | 201.116 | 0.96634 | 3.08E-16 | 2.24E-13 | TEC |
| ENSG00000222041 | CYTOR | 1848.989 | 1.391568 | 3.34E-16 | 2.38E-13 | lncRNA |
| ENSG00000272941 | ENSG00000272941 | 117.0442 | 1.174626 | 3.48E-16 | 2.42E-13 | lncRNA |
| ENSG00000120875 | DUSP4 | 291.3301 | 1.627276 | 4.13E-16 | 2.81E-13 | protein_coding |
| ENSG00000101365 | IDH3B | 624.1663 | -0.60318 | 5.20E-16 | 3.46E-13 | protein_coding |
| ENSG00000142871 | CCN1 | 536.5416 | 2.458004 | 6.21E-16 | 4.05E-13 | protein_coding |
| ENSG00000166527 | CLEC4D | 32.22037 | 3.859848 | 7.76E-16 | 4.95E-13 | protein_coding |

**Supplementary Table 2.**
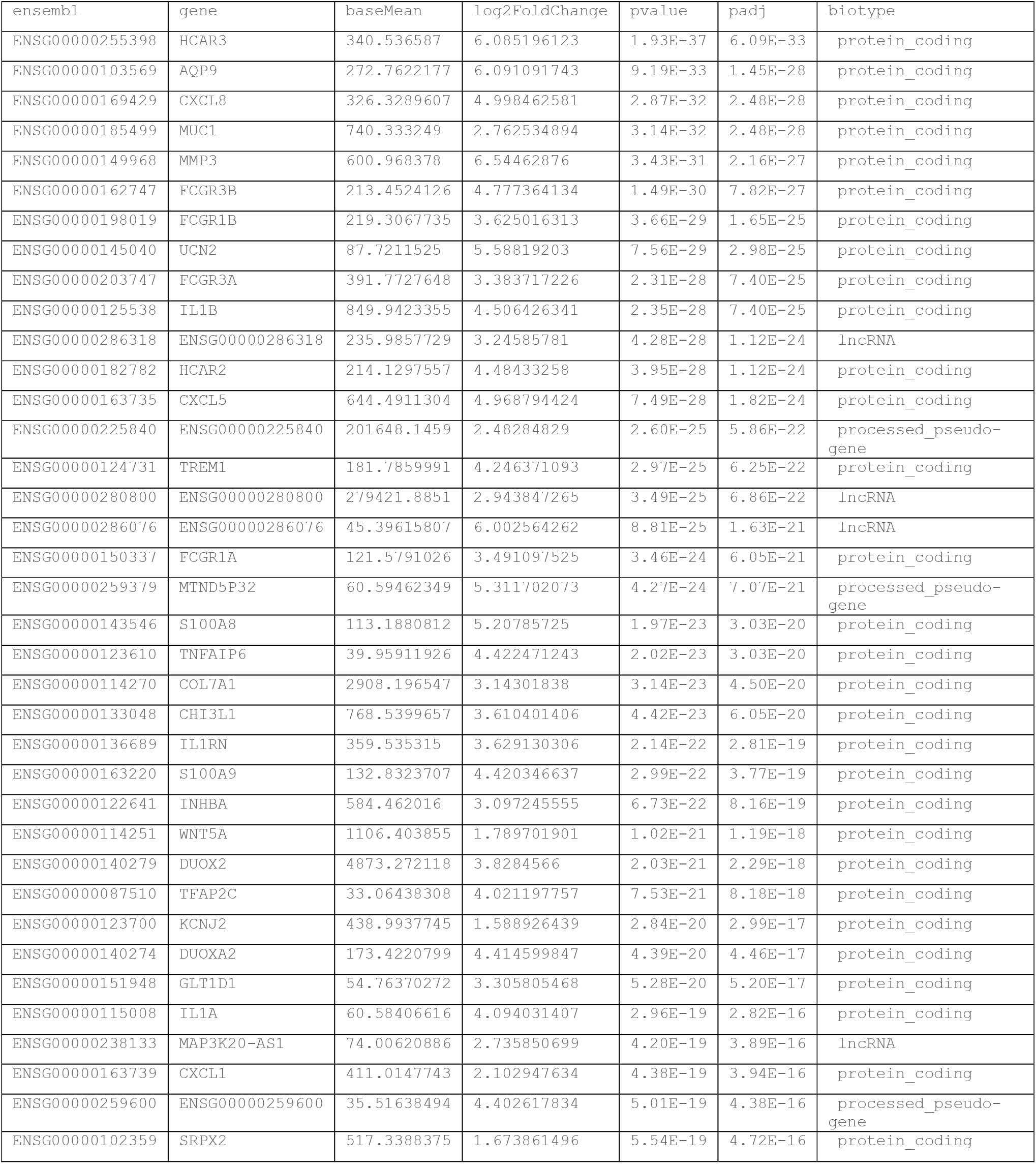
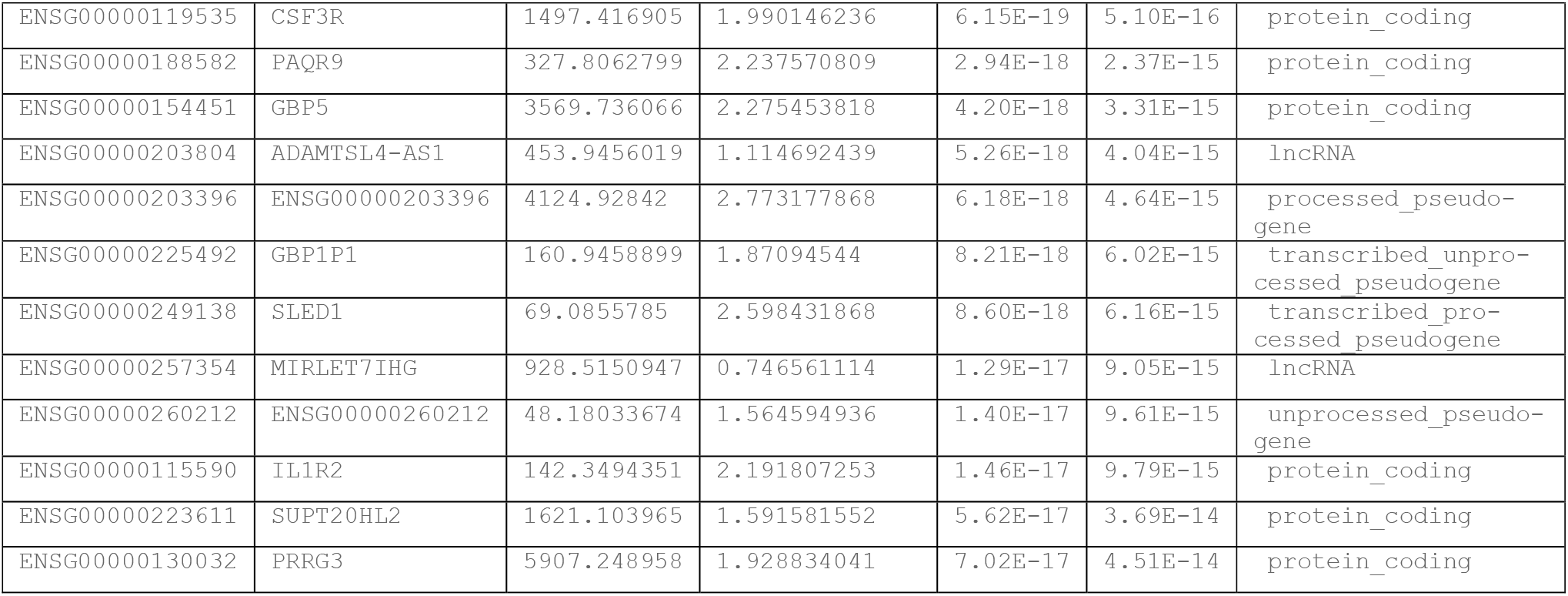

**Supplementary Table 3.**
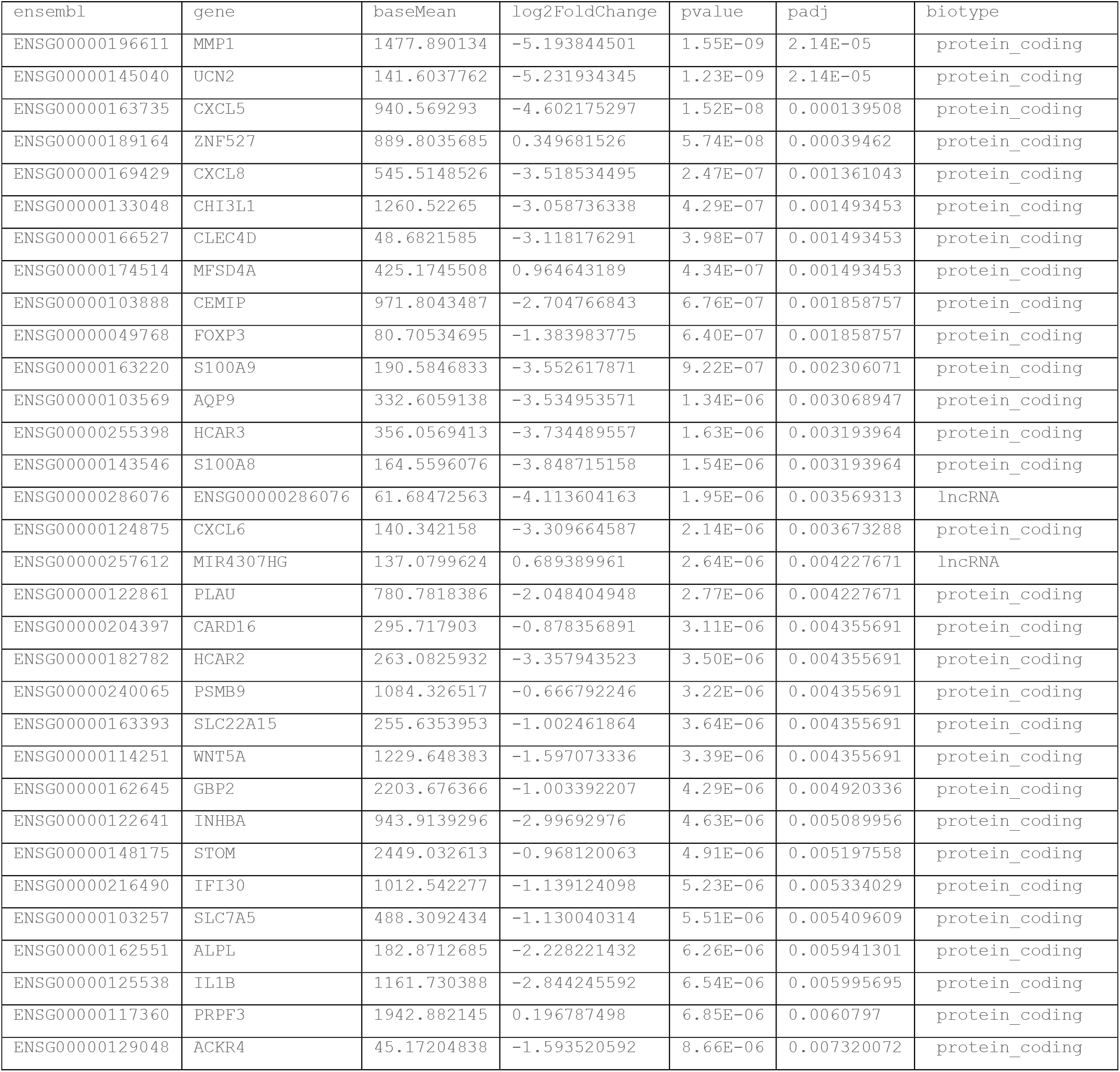
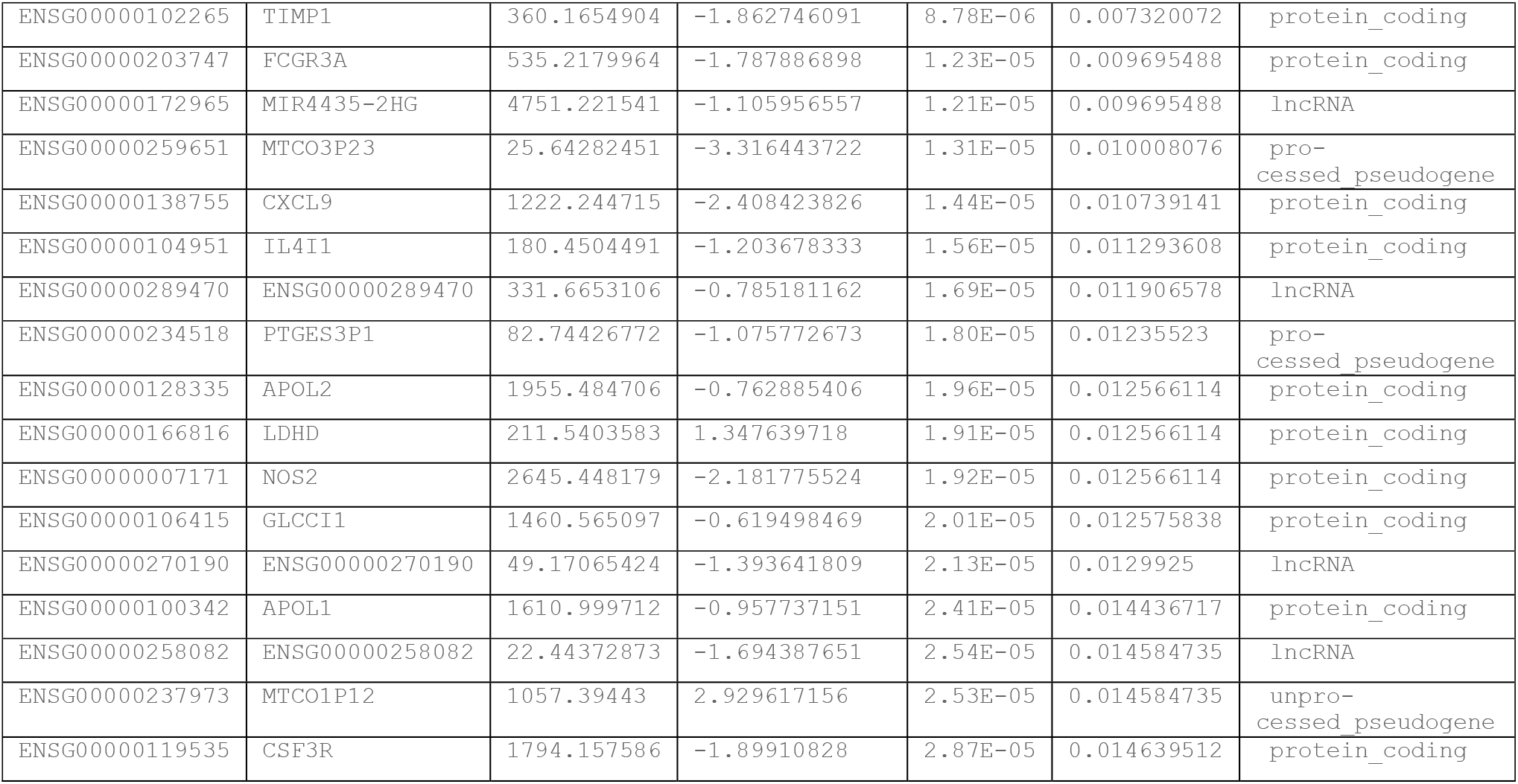

**Supplementary Table 4.**
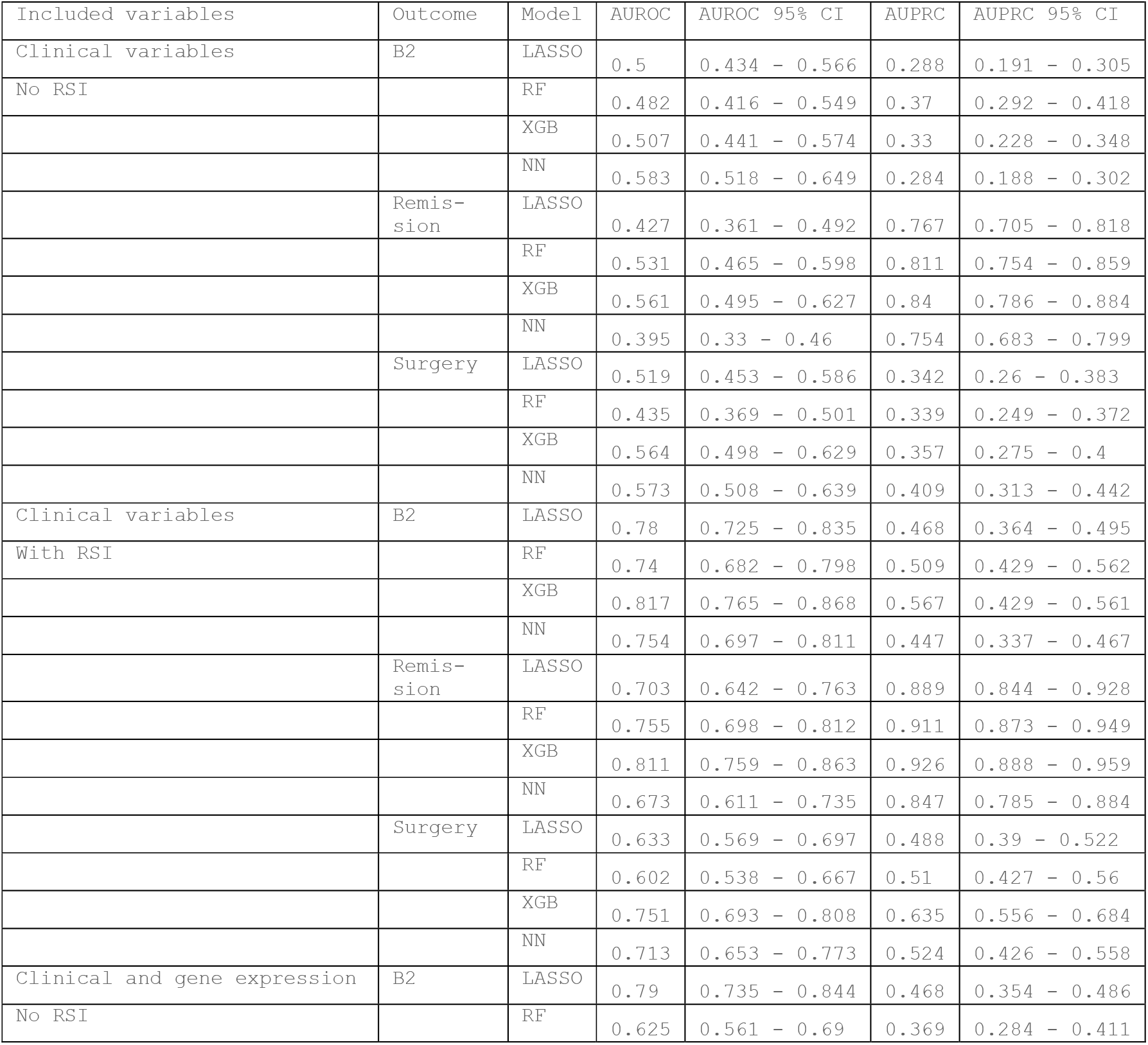
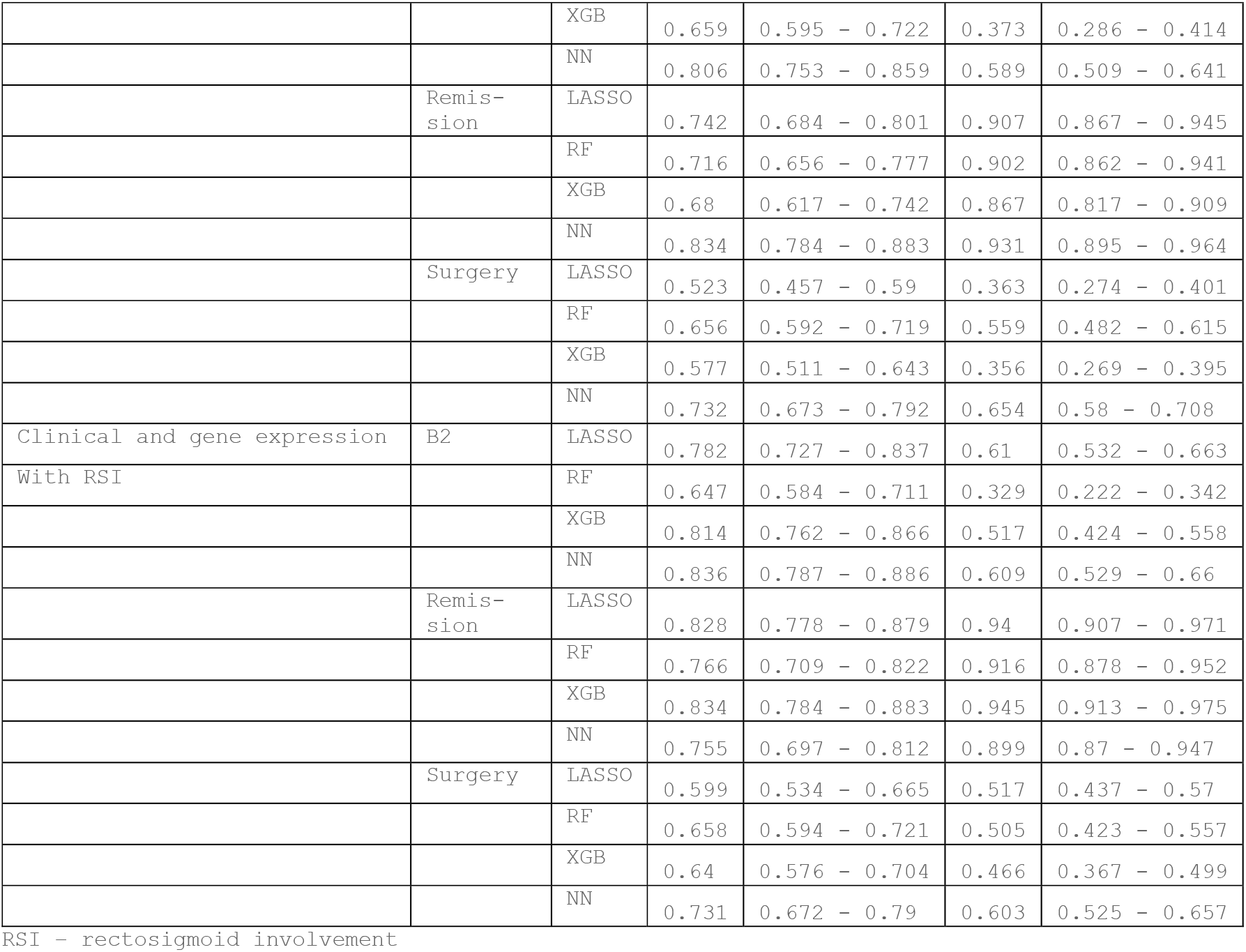

| Included variables | Outcome | Model | AUROC | AUROC 95% CI | AUPRC | AUPRC 95% CI |
| --- | --- | --- | --- | --- | --- | --- |
| Clinical variables | B2 | LASSO | 0.5 | 0.434 - 0.566 | 0.288 | 0.191 - 0.305 |
| No RSI |  | RF | 0.482 | 0.416 - 0.549 | 0.37 | 0.292 - 0.418 |
|  |  | XGB | 0.507 | 0.441 - 0.574 | 0.33 | 0.228 - 0.348 |
|  |  | NN | 0.583 | 0.518 - 0.649 | 0.284 | 0.188 - 0.302 |
|  | Remis-<br>sion | LASSO | 0.427 | 0.361 - 0.492 | 0.767 | 0.705 - 0.818 |
|  |  | RF | 0.531 | 0.465 - 0.598 | 0.811 | 0.754 - 0.859 |
|  |  | XGB | 0.561 | 0.495 - 0.627 | 0.84 | 0.786 - 0.884 |
|  |  | NN | 0.395 | 0.33 - 0.46 | 0.754 | 0.683 - 0.799 |
|  | Surgery | LASSO | 0.519 | 0.453 - 0.586 | 0.342 | 0.26 - 0.383 |
|  |  | RF | 0.435 | 0.369 - 0.501 | 0.339 | 0.249 - 0.372 |
|  |  | XGB | 0.564 | 0.498 - 0.629 | 0.357 | 0.275 - 0.4 |
|  |  | NN | 0.573 | 0.508 - 0.639 | 0.409 | 0.313 - 0.442 |
| Clinical variables | B2 | LASSO | 0.78 | 0.725 - 0.835 | 0.468 | 0.364 - 0.495 |
| With RSI |  | RF | 0.74 | 0.682 - 0.798 | 0.509 | 0.429 - 0.562 |
|  |  | XGB | 0.817 | 0.765 - 0.868 | 0.567 | 0.429 - 0.561 |
|  |  | NN | 0.754 | 0.697 - 0.811 | 0.447 | 0.337 - 0.467 |
|  | Remis-<br>sion | LASSO | 0.703 | 0.642 - 0.763 | 0.889 | 0.844 - 0.928 |
|  |  | RF | 0.755 | 0.698 - 0.812 | 0.911 | 0.873 - 0.949 |
|  |  | XGB | 0.811 | 0.759 - 0.863 | 0.926 | 0.888 - 0.959 |
|  |  | NN | 0.673 | 0.611 - 0.735 | 0.847 | 0.785 - 0.884 |
|  | Surgery | LASSO | 0.633 | 0.569 - 0.697 | 0.488 | 0.39 - 0.522 |
|  |  | RF | 0.602 | 0.538 - 0.667 | 0.51 | 0.427 - 0.56 |
|  |  | XGB | 0.751 | 0.693 - 0.808 | 0.635 | 0.556 - 0.684 |
|  |  | NN | 0.713 | 0.653 - 0.773 | 0.524 | 0.426 - 0.558 |
| Clinical and gene expression | B2 | LASSO | 0.79 | 0.735 - 0.844 | 0.468 | 0.354 - 0.486 |
| No RSI |  | RF | 0.625 | 0.561 - 0.69 | 0.369 | 0.284 - 0.411 |
|  |  | XGB | 0.659 | 0.595 - 0.722 | 0.373 | 0.286 - 0.414 |
|  |  | NN | 0.806 | 0.753 - 0.859 | 0.589 | 0.509 - 0.641 |
|  | Remis-sion | LASSO | 0.742 | 0.684 - 0.801 | 0.907 | 0.867 - 0.945 |
|  |  | RF | 0.716 | 0.656 - 0.777 | 0.902 | 0.862 - 0.941 |
|  |  | XGB | 0.68 | 0.617 - 0.742 | 0.867 | 0.817 - 0.909 |
|  |  | NN | 0.834 | 0.784 - 0.883 | 0.931 | 0.895 - 0.964 |
|  | Surgery | LASSO | 0.523 | 0.457 - 0.59 | 0.363 | 0.274 - 0.401 |
|  |  | RF | 0.656 | 0.592 - 0.719 | 0.559 | 0.482 - 0.615 |
|  |  | XGB | 0.577 | 0.511 - 0.643 | 0.356 | 0.269 - 0.395 |
|  |  | NN | 0.732 | 0.673 - 0.792 | 0.654 | 0.58 - 0.708 |
| Clinical and gene expression | B2 | LASSO | 0.782 | 0.727 - 0.837 | 0.61 | 0.532 - 0.663 |
| With RSI |  | RF | 0.647 | 0.584 - 0.711 | 0.329 | 0.222 - 0.342 |
|  |  | XGB | 0.814 | 0.762 - 0.866 | 0.517 | 0.424 - 0.558 |
|  |  | NN | 0.836 | 0.787 - 0.886 | 0.609 | 0.529 - 0.66 |
|  | Remis-sion | LASSO | 0.828 | 0.778 - 0.879 | 0.94 | 0.907 - 0.971 |
|  |  | RF | 0.766 | 0.709 - 0.822 | 0.916 | 0.878 - 0.952 |
|  |  | XGB | 0.834 | 0.784 - 0.883 | 0.945 | 0.913 - 0.975 |
|  |  | NN | 0.755 | 0.697 - 0.812 | 0.899 | 0.87 - 0.947 |
|  | Surgery | LASSO | 0.599 | 0.534 - 0.665 | 0.517 | 0.437 - 0.57 |
|  |  | RF | 0.658 | 0.594 - 0.721 | 0.505 | 0.423 - 0.557 |
|  |  | XGB | 0.64 | 0.576 - 0.704 | 0.466 | 0.367 - 0.499 |
|  |  | NN | 0.731 | 0.672 - 0.79 | 0.603 | 0.525 - 0.657 |

RSI - rectosigmoid involvement

## References

1. Kugathasan, S. & Hoffmann, R. The Incidence and Prevalence of Pediatric Inflammatory Bowel Disease (IBD) in the USA. J. Pediatr. Gastroenterol. Nutr. 39, S48– S49 (2004).

2. Benchimol, E. I. et al. Incidence, outcomes, and health services burden of very early onset inflammatory bowel disease. Gastroenterology 147, 803-813.e7 (2014).

3. Loftus, C. G. et al. Update on the incidence and prevalence of Crohn’s disease and ulcerative colitis in Olmsted County, Minnesota, 1940-2000. Inflamm. Bowel Dis. 13, 254–261 (2007).

4. Vernier-Massouille, G. et al. Natural History of Pediatric Crohn’s Disease: A Population-Based Cohort Study. Gastroenterology 135, 1106–1113 (2008).

5. Thia, K. T., Sandborn, W. J., Harmsen, W. S., Zinsmeister, A. R. & Loftus, E. V. Risk Factors Associated With Progression to Intestinal Complications of Crohn’s Disease in a Population-Based Cohort. Gastroenterology 139, 1147–1155 (2010).

6. Freeman, H. J. Age-dependent phenotypic clinical expression of Crohn’s disease. J. Clin. Gastroenterol. 39, 774–777 (2005).

7. Pigneur, B. et al. Natural history of Crohn’s disease: comparison between childhood- and adult-onset disease. Inflamm. Bowel Dis. 16, 953–961 (2010).

8. Abraham, B. P., Mehta, S. & El-Serag, H. B. Natural history of pediatric-onset inflammatory bowel disease: a systematic review. J. Clin. Gastroenterol. 46, 581–589 (2012).

9. Haberman, Y. et al. Pediatric Crohn disease patients exhibit specific ileal transcriptome and microbiome signature. J. Clin. Invest. 124, 3617–3633 (2014).

10. Neurath, M. F. Cytokines in inflammatory bowel disease. Nature Reviews Immunology vol. 14 329–342 (2014).

11. Noble, C. L. et al. Characterization of intestinal gene expression profiles in Crohn’s disease by genome-wide microarray analysis. Inflamm. Bowel Dis. 16, 1717–1728 (2010).

12. West, N. R. et al. Oncostatin M drives intestinal inflammation and predicts response to tumor necrosis factor-neutralizing therapy in patients with inflammatory bowel disease. Nat. Med. 23, 579–589 (2017).

13. Leal, R. F. et al. Identification of inflammatory mediators in patients with Crohn’s disease unresponsive to anti-TNFα therapy. Gut 64, 233–242 (2015).

14. Kugathasan, S. et al. Prediction of complicated disease course for children newly diagnosed with Crohn’s disease: a multicentre inception cohort study. Lancet 389, 1710–1718 (2017).

15. Haberman, Y. et al. Mucosal Inflammatory and Wound Healing Gene Programmes Reveal Targets for Stricturing Behaviour in Paediatric Crohn’s Disease. J. Crohn’s Colitis 15, 273–286 (2021).

16. Foster, J. D. et al. Application of objective clinical human reliability analysis (OCHRA) in assessment of technical performance in laparoscopic rectal cancer surgery. Tech. Coloproctol. 20, 361–367 (2016).

17. Isakov, O., Dotan, I. & Ben-Shachar, S. Machine Learning– Based Gene Prioritization Identifies Novel Candidate Risk Genes for Inflammatory Bowel Disease. Inflamm. Bowel Dis. 23, 1516–1523 (2017).

18. Ungaro, R. C. et al. Machine learning identifies novel blood protein predictors of penetrating and stricturing complications in newly diagnosed paediatric Crohn’s disease. Aliment. Pharmacol. Ther. 53, 281–290 (2021).

19. Gardiner, L. J. et al. Combining explainable machine learning, demographic and multi-omic data to inform precision medicine strategies for inflammatory bowel disease. PLoS One 17, e0263248 (2022).

20. Kugathasan, S. et al. Prediction of complicated disease course for children newly diagnosed with Crohn’s disease: a multicentre inception cohort study. Lancet 389, 1710–1718 (2017).

21. Keith, B. P. et al. Colonic epithelial miR-31 associates with the development of Crohn’s phenotypes. JCI insight 3, (2018).

22. Satsangi, J., Silverberg, M. S., Vermeire, S. & Colombel, J. F. The Montreal classification of inflammatory bowel disease: controversies, consensus, and implications. Gut 55, 749–753 (2006).

23. Patro, R., Duggal, G., Love, M. I., Irizarry, R. A. & Kingsford, C. Salmon provides fast and bias-aware quantification of transcript expression. Nat. Methods 14, 417–419 (2017).

24. Wang, L. et al. Measure transcript integrity using RNA-seq data. BMC Bioinformatics 17, 1–16 (2016).

25. Risso, D., Ngai, J., Speed, T. P. & Dudoit, S. Normalization of RNA-seq data using factor analysis of control genes or samples. Nat. Biotechnol. 32, 896–902 (2014).

26. Love, M. I., Huber, W. & Anders, S. Moderated estimation of fold change and dispersion for RNA-seq data with DESeq2. Genome Biol. 15, 1–21 (2014).

27. Ritchie, M. E. et al. limma powers differential expression analyses for RNA-sequencing and microarray studies. Nucleic Acids Res. 43, e47–e47 (2015).

28. Robinson, M. D., McCarthy, D. J. & Smyth, G. K. edgeR: a Bioconductor package for differential expression analysis of digital gene expression data. Bioinformatics 26, 139–140 (2010).

29. Sergushichev, A. A. An algorithm for fast preranked gene set enrichment analysis using cumulative statistic calculation. bioRxiv 060012 (2016) doi:10.1101/060012.

30. Liberzon, A. et al. The Molecular Signatures Database (MSigDB) hallmark gene set collection. Cell Syst. 1, 417 (2015).

31. Blighe, K., Rana, S. & Lewis, M. EnhancedVolcano: Publication-ready volcano plots with enhanced colouring and labeling. R package version 1.14.0. (2022).

32. R Core Team. R: A Language and Environment for Statistical Computing. (2020).

33. Levine, A. et al. Complicated Disease and Response to Initial Therapy Predicts Early Surgery in Paediatric Crohn’s Disease: Results From the Porto Group GROWTH Study. J. Crohn’s Colitis 14, 71–78 (2020).

34. Géron, A. Hands-on machine learning with Scikit-Learn, Keras, and TensorFlow: Concepts, tools, and techniques to build intelligent systems. (O’Reilly Media, 2019).

35. Pedregosa, F. et al. Scikit-learn: Machine Learning in Python. J. Mach. Learn. Res. 12, 2825–2830 (2011).

36. scikit learn. https://scikit-learn.org/stable/modules/generated/sklearn.linear_model.LogisticRegression.html.

37. Chollet, F. & others. Keras. (2015).

38. Alfredsson, J. & Wick, M. J. Mechanism of fibrosis and stricture formation in Crohn’s disease. Scand. J. Immunol. 92, e12990 (2020).

39. Ta, A. D. et al. Association of Baseline Luminal Narrowing With Ileal Microbial Shifts and Gene Expression Programs and Subsequent Transmural Healing in Pediatric Crohn Disease. Inflamm. Bowel Dis. 27, 1707– 1718 (2021).

40. Toyonaga, T. et al. Increased colonic expression of ACE2 associates with poor prognosis in Crohn’s disease. Sci. Rep. 11, (2021).

41. Kofla-Dlubacz, A., Matusiewicz, M., Krzystek-Korpacka, M. & Iwanczak, B. Correlation of MMP-3 and MMP-9 with Crohn’s Disease Activity in Children. Dig. Dis. Sci. 57, 706 (2012).

42. Van Beelen Granlund, A. et al. REG gene expression in inflamed and healthy colon mucosa explored by in situ hybridisation. Cell Tissue Res. 352, 639 (2013).

43. Torres, J. et al. Predicting Outcomes to Optimize Disease Management in Inflammatory Bowel Diseases. J. Crohn’s Colitis 10, 1385–1394 (2016).

44. Waljee, A. K. et al. Predicting Hospitalization and Outpatient Corticosteroid Use in Inflammatory Bowel Disease Patients Using Machine Learning. Inflamm. Bowel Dis. 24, 45 (2018).

45. Waljee, A. K. et al. Development and Validation of Machine Learning Models in Prediction of Remission in Patients With Moderate to Severe Crohn Disease. JAMA Netw. Open 2, (2019).

46. Kc, C. et al. Predicting Risk of Postoperative Disease Recurrence in Crohn’s Disease: Patients With Indolent Crohn’s Disease Have Distinct Whole Transcriptome Profiles at the Time of First Surgery. Inflamm. Bowel Dis. 25, 180–193 (2019).

47. Ungaro, R. C. et al. Machine learning identifies novel blood protein predictors of penetrating and stricturing complications in newly diagnosed paediatric Crohn’s disease. Aliment. Pharmacol. Ther. 53, 281–290 (2021).

48. Noor, N. M., Sousa, P., Paul, S. & Roblin, X. Early Diagnosis, Early Stratification, and Early Intervention to Deliver Precision Medicine in IBD. Inflamm. Bowel Dis. 28, 1254– 1264 (2022).

49. Gubatan, J. et al. Artificial intelligence applications in inflammatory bowel disease: Emerging technologies and future directions. World J. Gastroenterol. 27, 1920–1935 (2021).

